# Construction and Functional Characterization of a Heterologous Quorum Sensing Circuit in *Clostridium sporogenes*

**DOI:** 10.1101/2025.08.21.671393

**Authors:** Sara Sadr, Bahram Zargar, Marc G Aucoin, Brian Ingalls

## Abstract

Quorum sensing (QS) is a bacterial communication mechanism that regulates gene expression in a population density-dependent manner. Substantial progress has been made in the engineering of QS systems in Gram-negative bacteria, but development of engineered QS systems in Gram positive bacteria remains limited. In this study, *obligate anaerobic Gram-positive bacterium Clostridium sporogenes* was engineered with the *Staphylococcus aureus* agr-QS system to enable density-dependent gene regulation. Using LC-MS/MS, we confirmed production of autoinducing peptides in the engineered *C. sporogenes* strain. A QS-regulated GFP reporter demonstrated activation of expression in response to both exogenous AIP addition and increasing cell density, confirming functional integration of the agr operon. A media refreshment experiment showed that replacing the culture supernatant delayed QS activation, highlighting the importance of signal accumulation. Moreover, we observed that a non-cognate AIP from another agr specificity group acts as a competitive antagonist, inhibiting gene expression under the QS promoter. To our knowledge, this study presents the first successful engineering of the agr quorum sensing system in an obligate anaerobe, expanding the synthetic biology toolkit and offers new opportunities for bacterial therapies and metabolic engineering.

## Introduction

Quorum sensing (QS) is a communication system through which bacteria regulate gene expression based on population density (Waters & Bassler, 2005; Whiteley & Stephen, 2017). By sensing the concentration of small signaling molecules, bacteria coordinate multicellular behaviors such as bioluminescence (Bassler, Wright, & Stiverman, 1994; Engebrecht, Nealson, & Silverman, 1983; Verma & Miyashiro, 2013), biofilm formation (Hammer & Bassler, 2003; Li & Zhao, 2020), secondary metabolite production (Barnard et al., 2007; Pena et al., 2019), and virulence (Jenul & Horswill, 2019; Rutherford & Bassler, 2012). QS is best understood in Gram-negative bacteria, in which it typically involves acyl-homoserine lactones (AHLs) as signaling molecules that diffuse freely across cell membranes. As the bacterial population grows, the concentration of AHLs increases. AHLs bind to specific intracellular receptor proteins, such as LuxR-type transcriptional regulators, activating QS-regulated gene expression and increasing AHL production in a positive feedback loop. In contrast, Gram-positive bacteria rely on actively secreted small autoinducing peptides (AIPs). AIPs are detected by specific membrane-bound proteins, which are typically part of a two-component regulatory system composed of a sensor histidine kinase and a response regulator (Kleerebezem & Quadri, 1997; Novick & Geisinger, 2008). Upon AIP binding to its receptor, the sensor kinase undergoes autophosphorylation, initiating a signaling cascade that phosphorylates the response regulator. This activates or represses QS-regulated genes, inducing the QS-response and triggering increased AIP secretion in a positive feedback loop (Novick & Geisinger, 2008).

Quorum sensing (QS) is widely used in synthetic biology to engineer bacterial populations with predictable and programmable behavior. QS systems have been integrated into circuits to enable responses to environmental signals or population density, facilitating advancements in biotechnology and biomanufacturing (Alnahhas et al., 2020; Danino, Mondragón-Palomino, Tsimring, & Hasty, 2010; Dinh, Chen, & Prather, 2020; Fedorec, Karkaria, Sulu, & Barnes, 2021; Gu et al., 2020; Gupta, Reizman, Reisch, & Prather, 2017; E. M. Kim et al., 2017; J. K. Kim et al., 2019). Exploiting QS has emerged as a valuable strategy for therapeutic applications, particularly for controlling gene expression and microbial populations in disease contexts. For example, as a tumor-targeting therapy, a *Salmonella sp.* was engineered to express a therapeutic gene in a tumor only when the density of *Salmonella* reaches a threshold (Swofford, Van Dessel, & Forbes, 2015). QS-controlled lysis circuits engineered in *Escherichia coli* enable pulsatile release of therapeutic agents for targeted drug delivery in tumors (Din et al., 2016), or the production of nanobodies (anti-CD47) and their release in tumor environments, the latter demonstrating effective tumor regression *in vitro* and *in vivo* (Chowdhury et al., 2019). QS-based strategies have also been applied to infection control. Quorum sensing inhibitors (QSIs) have been explored to disrupt bacterial communication and biofilm formation, addressing antibiotic resistance (Ellermann & Sperandio, 2021). Additionally, QS-regulated systems are being developed for therapeutic bacteria to treat intestinal diseases (Dang et al., 2023). These advancements underscore the potential of QS in improving precision and efficacy in medical applications.

While QS systems in Gram-negative bacteria are frequently studied and widely applied in synthetic biology, the engineering and application of QS in Gram-positive bacteria remains less explored. A well-characterized QS system in Gram-positive bacteria is found in *Staphylococcus aureus*, where it regulates pathogenicity and adaptability by controlling virulence factor production and biofilm formation (Novick & Geisinger, 2008; Yamazaki, Ito, Tamai, Nakagawa, & Nakamura, 2024; Yarwood, Bartels, Volper, & Greenberg, 2004). Central to this system is the accessory gene regulator (agr) locus, which functions as a key regulatory circuit for bacterial communication (Figure 1). The agr system produces and detects autoinducing peptides (AIPs) that accumulate in the environment as the bacterial population grows. Extracellular AIPs interact with the AgrC receptor, a membrane-bound sensor kinase, triggering a phosphorylation cascade via the response regulator AgrA. This cascade activates the transcription of downstream genes, including RNAIII, a key effector molecule that modulates the expression of virulence factors, such as toxins, enzymes, and surface proteins (Bronesky et al., 2016). To date, only a few studies have reported the successful engineering of synthetic QS systems in Gram-positive hosts. The earliest such study was conducted by Marchand and Collins, who introduced a modified version of the agr system from *Staphylococcus aureus* into the Gram-positive host *Bacillus megaterium* to create a functional synthetic QS circuit, using a xylose-inducible promoter to control *agrBDCA* expression (Marchand & Collins, 2016). These efforts established the first synthetic peptide-based QS system in a Gram-positive host and highlighted autoinduction, dynamic regulation and metabolic engineering. More recently, a synthetic QS system based on LuxR-type regulators and homoserine lactones was engineered in the Gram-positive host *Bacillus subtilis*, demonstrating that non-native QS systems can function in Gram-positive bacteria when equipped with appropriate regulatory elements (Zeng, Sarker, Howitz, Shah, & Andrews, 2024). This further highlights the expanding potential of Gram-positive hosts for synthetic QS applications.

**Figure 1.**
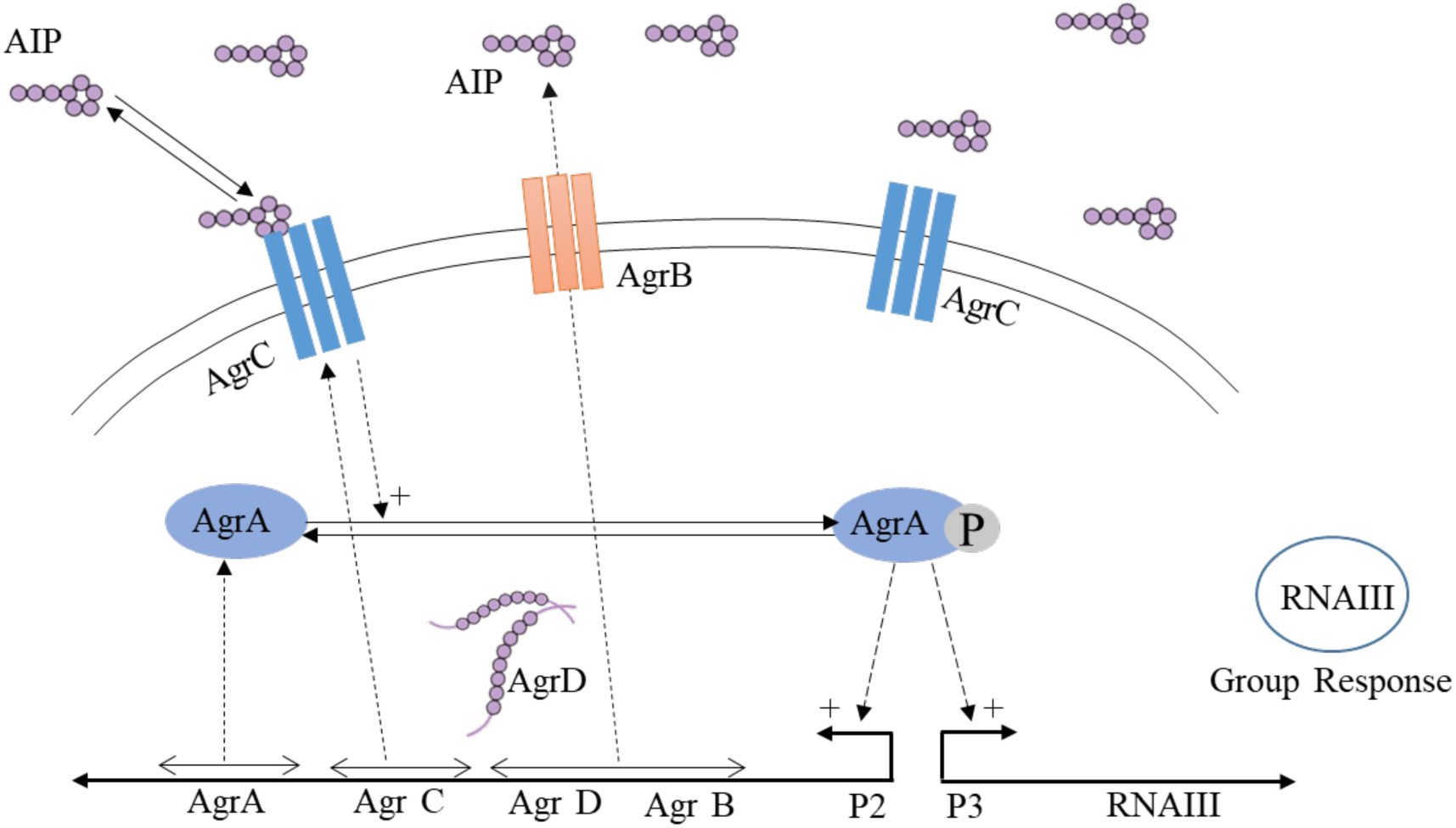
Schematic representation of the quorum sensing (QS) Agr system in S*taphylococcus aureus.* The Agr system includes the production of autoinducing peptides (AIP) by AgrD, which are processed and exported by AgrB. As the AIP concentration increases, AIP binds to the receptor AgrC, triggering a phosphorylation cascade that activates AgrA. Activated AgrA induces the expression of the Agr operon and of RNAIII, which regulates virulence factors and coordinates group behaviors.

*Clostridium sporogenes* is a non-pathogenic, obligate anaerobic, and spore-forming bacterium that has garnered significant interest for its unique biological properties (Kubiak et al., 2015). It has been extensively studied for its potential in treating solid-mass tumors. Spores of *Clostridium sp.* remain inert in normoxic healthy tissue but germinate in the hypoxic regions of solid tumors, enabling precise targeting of necrotic cores and high levels of tumor colonization compared to other *Clostridium* species (Fox et al., 1996; Lemmon et al., 1997; Liu, Minton, Giaccia, & Brown, 2002; Mowday et al., 2016). Beyond its therapeutic potential, *C. sporogenes* has also been investigated for biofuel production (Gottumukkala et al., 2013, 2015).

Leveraging the coordinated regulatory mechanism of the agr system of *S. aureus*, we engineered its operon into *C. sporogenes*. We confirmed the system’s selective response to the *S. aureus* signaling peptide AIP-III and confirmed that at high population density the engineered strain produces AIP-III. We then confirmed density-dependent activation in batch growth and that removal of accumulated AIP-III by supernatant refreshment disrupts the density-dependent timing of activation during growth. In addition, we observed that a non-cognate AIP from another agr specificity groups acts as a competitive antagonist, inhibiting gene expression driven by the QS promoter. This engineered system enables QS-driven control of gene expression, achieving coordinated population density-dependent behavior, providing a path to therapeutic actions initiated only above a population density threshold within target tissues as in (Swofford et al., 2015).

## Materials and Methods

### Bacteria strains and plasmids

*C. sporogenes* NCIMB 10696 (referred to here as the Native strain) and *E. coli* CA434 were generously provided by Professor Nigel Minton (University of Nottingham, UK) and Professor Mike Young (Aberystwyth University, UK), respectively. The shuttle vector pMTL8225x (Heap, Pennington, Cartman, & Minton, 2009) was also provided by Professor Minton. Genomic DNA from *S. aureus* (ATCC 43300) was obtained from the American Type Culture Collection (VA, USA). All restriction enzymes were sourced from New England Biolabs (Whitby, ON, Canada). The plasmid pGlow-Xn-Pp1-CI was purchased from BioCat GmbH (Heidelberg, Germany).

Plasmids pAG3, encoding the agr operon and GFP expression driven by the P3 promoter (which in its native context controls transcription of the regulatory RNAIII (Figure 1)), and pTG, coding for constitutive GFP expression (to be used as a control), were constructed using the pMTL8225x shuttle vector as the backbone.

To construct pAG3, first, the evoglow gene was amplified from the pGlow-Xn-Pp1-CI plasmid using pglow_f_XbaI and pglow_r_pstI_spei primers (see Supplementary Table S1 for all primer sequences), and the resulting PCR product was cloned into the XbaI site of pMTL8225x to create pMTL_evoglow, which was subsequently linearized with XbaI. The agr locus was amplified from *S. aureus* ATCC 43300 genomic DNA using Agr43300_fwd and Agr43300_rev primers. Both the linearized pMTL_evoglow vector and the amplified agr locus insert were separated by gel electrophoresis, and the specific bands (∼5.8 kb for the vector and ∼3 kb for the insert) were extracted and purified using GenepHlow Gel/PCR Kit (DFH100, Geneaid Biotech Ltd., New Taipei City, Taiwan). The vector and insert were then amplified using Gibson primers and assembled using the HiFi Gibson Assembly Kit (New England Biolabs, Ipswich, MA, USA).

To construct pTG, the constitutive thl promoter and the evoglow gene from pGlow-Xn-Pp1-CI were PCR amplified with the thl_F and pGlow_PstI_SpeI primers, respectively, and inserted into the EcoRI-XbaI site of pMTL8225x.

Plasmid maps for both pAG3 and pTG are provided in the Supplement, Figure S1. The plasmids pAG3 and pTG were transformed into competent *E. coli* CA434 cells by electroporation using a Gene Pulser Xcell Electroporation System (Bio-Rad Laboratories Ltd, Mississauga, ON, Canada). The preset protocol for *E. coli* was used with a 2 mm cuvette. The plasmids were transferred into *C. sporogenes* by conjugation following the procedure described by Theys et al. (Theys et al., 2006). The transformed donor *E. coli* was inoculated in LB broth with erythromycin and incubated overnight at 37 °C in an orbital shaker at 225 rpm. The target *C. sporogenes* strain was grown in TYG medium under anaerobic conditions overnight at 37 °C. On the second day, *C. sporogenes* was subcultured, and 1 mL of the overnight *E. coli* donor culture was pelleted by centrifugation at 8000 rpm for 1 min, then washed twice with 0.5 mL sterile PBS. The *E. coli* pellet was re-suspended in 200 µL of late exponential-phase *C. sporogenes* culture to create a conjugation mixture, which was then spotted onto a non-selective TYGA plate (3% trypticase, 2% yeast extract, 0.1% glucose, and 1.5% bacterial agar). The plate was incubated at 37 °C in anaerobic conditions for 16 h to allow plasmid transfer. Following conjugation, 1 mL of anaerobic sterile PBS was added to the plate, and the cell layer was scraped and re-suspended in PBS. The cell-PBS slurry was transferred to a micro-tube, and 100 µL of the slurry and its 10-fold dilutions were spread onto TYGA plates containing 250 µg mL⁻¹ D-cycloserine (to select against *E. coli*) and 10 µg mL⁻¹ erythromycin (to select for the plasmid). After 48 h at 37 °C, colonies were selected for further study, with *C. sporogenes* strains containing plasmids pAG3 or pTG, which were then referred to as PAG3 and PTG, respectively.

To confirm the resultant transconjugants, plasmids were extracted using the Invitrogen PureLink Quick Plasmid Miniprep Kit (Invitrogen, Thermo Fisher Scientific, Waltham, MA, USA) with slight modifications. Bacterial cultures were harvested during the exponential phase at an OD_600_ of 0.9. Cells were washed twice with TSE buffer (50 mM Tris-HCl, 25 mM EDTA, 6.7% sucrose, pH 8), and the pellet was resuspended in 250 µL of resuspension buffer (50 mM Tris-HCl, 10 mM EDTA, pH 8) supplemented with 0.1 mg mL⁻¹ RNase A. Then, 250 µL of lysis buffer with 47 µL of freshly prepared 100 mg mL⁻¹ lysozyme solution (Sigma, 62970) was added, and the mixture was gently inverted. Next, 300 µL of TE+SDS buffer (50 mM Tris-HCl, 20 mM EDTA, 4% SDS, pH 8) was added and gently mixed by inversion, followed by incubation at 37°C for 2 h. The subsequent steps, including binding, washing, and elution of plasmid DNA, were performed as per the kit instructions. Plasmid samples were verified by Full-length plasmid sequencing using Oxford Nanopore Technology (Plasmidsaurus).

### Bacterial Growth Conditions

Anaerobic growth of *Clostridium* strains was conducted in a Thermo Forma Anaerobic System (Model 1025) under an atmosphere of 5% carbon dioxide (CO₂), 10% hydrogen (H₂), and 85% nitrogen (N₂) at 37 °C, using TYG medium (3% trypticase, 2% yeast extract, and 0.1% glucose). *E. coli* was grown in LB broth (1% trypticase, 0.5% yeast extract, and 1% NaCl) or on LB agar (1.5% agar) at 37 °C. For transformed *E. coli* strains, erythromycin was added at a concentration of 500 µg mL⁻¹, while C. *sporogenes* cultures were supplemented with 10 µg mL⁻¹ erythromycin. Both *E. coli* and *C. sporogenes* cultures were stored at −80 °C in glass cryovials containing 15% v/v and 10% v/v glycerol, respectively.

### Growth and Fluorescence Assay

Bacterial cultures were grown in 50 mL tubes in an anaerobic chamber with a total culture volume of 30 mL in TYG medium. Samples (1 mL) were collected at various time points following the start of exponential growth. Each sample was washed twice in 1 mL phosphate-buffered saline (PBS) before measuring optical density (OD_600_) and the intensity of green fluorescent protein (GFP) using a Cytation 5 plate reader (BioTek). In the media replacement protocol, bacteria cultures were centrifuged after 7 hours at 10,000 x g for 10 minutes. The supernatant was discarded, and fresh TYG medium (pre-warmed to 37 °C and supplemented with erythromycin) was added.

To evaluate the response of the synthetic QS system to external signaling molecules, AIP-III (SB Peptides, France, catalog number: CRB1001688) was added to the culture medium at a final concentration of 1 μM. AIP was added at the time of inoculation, and cultures were incubated under anaerobic conditions in TYG medium. Samples were collected at multiple time points, as described above, to monitor bacterial growth and GFP expression. Assessment of the response to AIP-I (Biosynthetech, catalog number: CRB1000249) followed the same protocol, as did the negative control with no AIP addition.

### Liquid Chromatography–Mass Spectrometry (LC-MS) Analysis of AIP

The Native and PAG3 strains were cultured in TYG medium under anaerobic conditions at 37 °C. After 10 hours of incubation, cultures were centrifuged at 10,000 × g for 10 minutes at 4 °C. The resulting supernatant was collected and stored on ice prior to peptide extraction.

Solid-phase extraction (SPE) was performed using Strata® C18-E SPE cartridges (200 mg / 3 mL, 55 µm, 70 Å pore size; Phenomenex, 8B-S001-FBJ) according to the manufacturer’s instructions. Briefly, cartridges were activated with 3 mL methanol and equilibrated with 3 mL Milli-Q water. A total of 10 mL of culture supernatant was loaded per cartridge, followed by washing with 3 mL water. Bound peptides were eluted with 3 mL of 80% acetonitrile (ACN) in water containing 0.1% formic acid.

LC-MS analysis was carried out using an API 3000 triple quadrupole mass spectrometer (Applied Biosystems/MDS SCIEX) coupled to an Agilent 1100 Series HPLC system (Hewlett-Packard). Chromatographic separation was performed using a Waters XBridge™ C18 column (3.5 µm, 4.6 × 100 mm; Part No. 186003033) maintained at 40 °C. An isocratic elution program was used with a mobile phase consisting of water with 0.1% formic acid (solvent A) and methanol with 0.1% formic acid (solvent B) in a 30:70 (v/v) ratio. The total run time was 10 minutes at a flow rate of 0.3 mL/min.

A multiple reaction monitoring (MRM) method was developed to selectively detect the predicted autoinducing peptide (AIP). The MRM transitions were optimized using a synthetic AIP-III standard peptide (sequence: INCDFLL), purchased from SB Peptides (France), catalog number CRB1001688. A 10 µM working solution of the standard was prepared in water and used for method development and calibration. Ionization was performed in electrospray ionization in positive ion mode (ESI+). Multiple Reaction Monitoring (MRM) transitions were optimized by direct infusion of the standard into the mass spectrometer. The following transitions were selected for quantification and validation: T1: 819.300 → 706.400 Da, T2: 819.300 → 592.200 Da, and T3: 819.300 → 479.300 Da. Optimized instrument parameters for each transition were: Declustering Potential (DP): 34.5 V, Entrance Potential (EP): 8.5 V, Collision Energy (CE): 36.5 V, and Collision Cell Exit Potential (CXP): 21.0 V. Data acquisition and analysis were performed using Analyst™ software (SCIEX). Quantification of AIP in culture supernatants was based on peak area integration of the AIP-specific MRM signal.

## Results

### Engineered PAG3 strain produces *S. aureus* agr AIP-III

After construction of the PAG3 strain, we assessed culture supernatant using LC-MS/MS in MRM mode to detect the presence of AIP-III, which is the AIP employed by the *S. aureus* agr system introduced into PAG3. A synthetic AIP-III standard (10 µM, 20 µL injection) was employed as a reference, and the supernatant from the Native strain and blank media were analyzed as negative controls. The standard yielded two distinct peaks, at retention times 5.72 min and 6.28 min, indicating stereoisomer formation during peptide synthesis (Figure 2A). The peak at 5.72 min co-eluted with a compound in the PAG3 culture supernatant (Figure 2B), confirming the presence of AIP-III. Quantification was performed based on T1 transition (T1: 819.3 → 706.4 Da), with 56% of the total signal in the standard corresponding to the biologically relevant isomer, equivalent to 111.8 pmol. The concentration of AIP-III in the PAG3 supernatant was calculated as 534 pmol/mL. No AIP signal was detected in the samples from the blank media or from cultures of the Native strain (Figure 2C, D).

**Figure 2.**
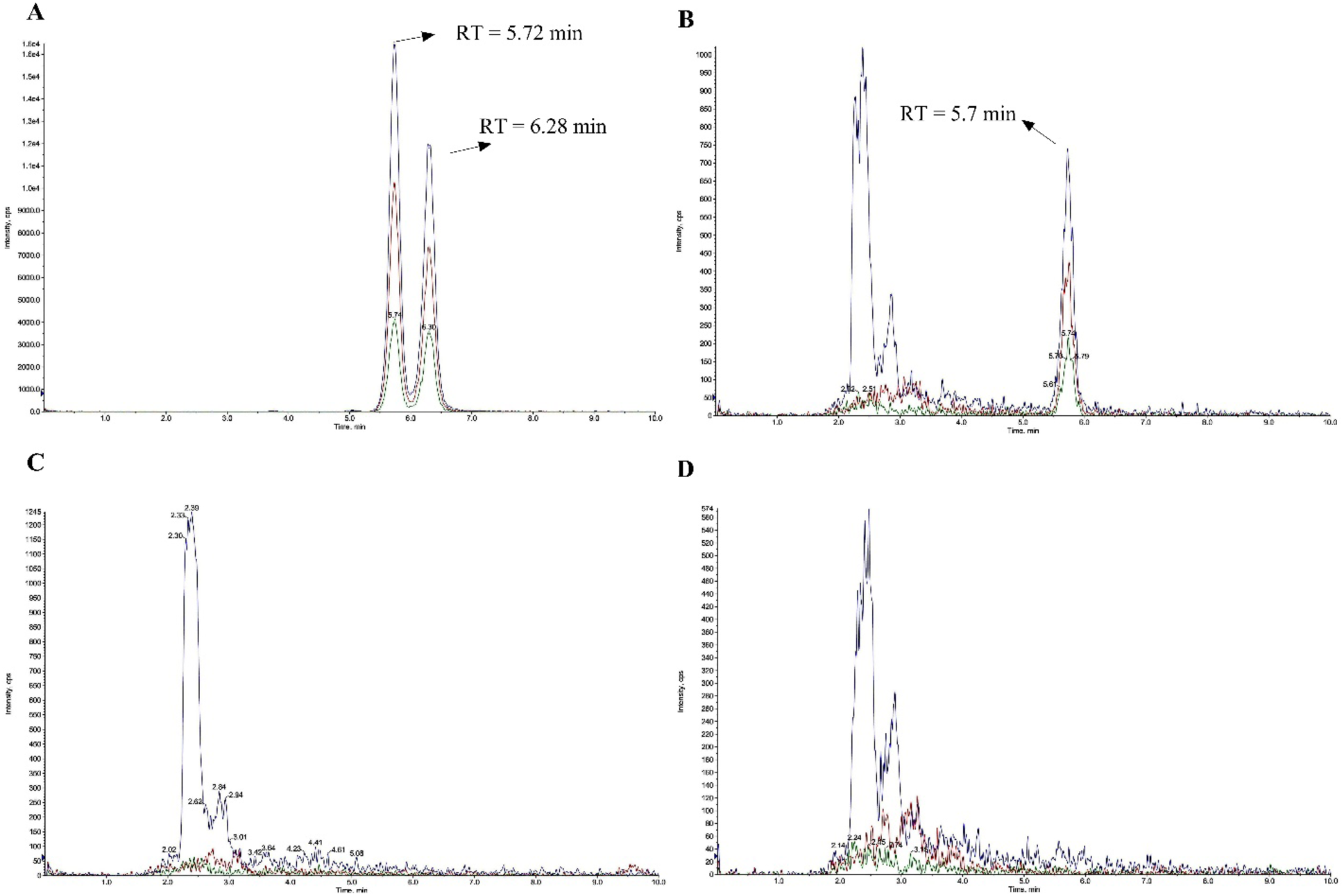
LC-MS/MS chromatograms of synthetic and sample-derived AIP-III detected by LC-MS/MS. A) Synthetic AIP-III standard (10 µM, 20 µl injection) showing two peaks at 5.72 and 6.28 min, corresponding to stereoisomers. B) Supernatant from the engineered PAG3 strain showing a single peak at 5.7 min corresponding to the biologically relevant isomer. C) Blank media showing no detectable signal at the AIP retention time. D) Supernatant sample from Native *C. sporogenes* showing no detectable AIP-III signal. Chromatograms display MRM transitions: blue (T1, 819.3 → 706.4 Da), red (819.3 → 592.2 Da), and green (819.3 → 479.3 Da). Quantification was based on the T1 transition and the peak at 5.7 min.

### Engineered agr operon responds selectively to AIP-III

To investigate the behavior of the synthetic QS system in *C. sporogenes*, PAG3 and Native strains were cultured in TYG medium, and 1µM of AIP-III was added to the culture at the time of inoculation. As a control, separate cultures of PAG3 and Native strains were grown without AIP-III supplementation. All cultures were grown in anaerobic conditions. Figure 3A shows the bacterial growth profiles over time, measured by OD_600_. The growth curves follow a similar trend, with no major difference in overall growth dynamics between supplemented and unsupplemented cultures. Figure 3B presents the fluorescence intensity measurements over the course of experiment. In the PAG3 strain, the fluorescence intensity increases more sharply over time in the presence of AIP-III compared to the unsupplemented control. In contrast, for the Native strain, the addition of AIP-III had no effect on fluorescence.

**Figure 3.**
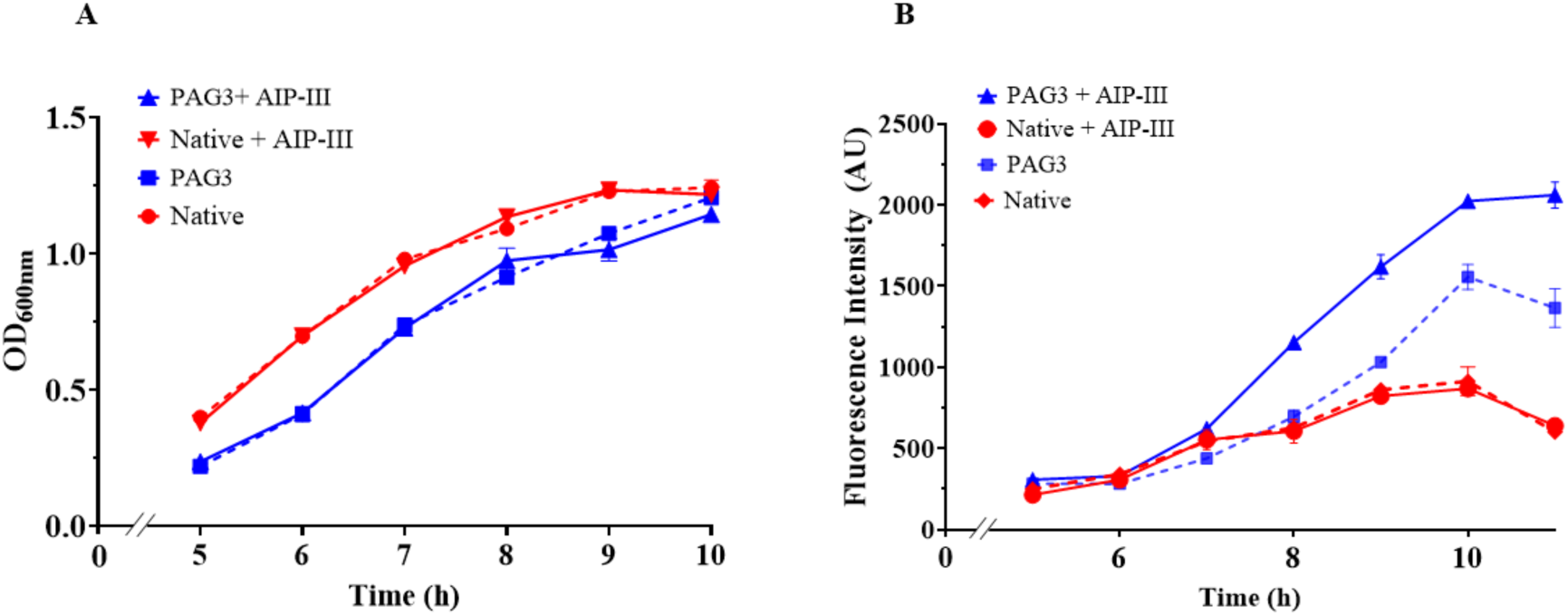
Growth and fluorescence intensity of PAG3 and Native strains with (solid lines) and without (dashed lines) exogenous AIP-III supplementation. A) OD_600_ growth curves of Native and PAG3 strains with and without AIP-III supplementation, B) Fluorescence intensity over time. Supplementation of AIP-III results in a marked increase in fluorescence in PAG3, compared to the unsupplemented PAG3 culture. The Native strain shows similar fluorescence levels in both conditions. Error bars represent one standard deviation from triplicate cultures.

To assess the QS system’s response to the addition of a non-cognate AIP signal, PAG3 and Native strains were cultured in TYG medium supplemented with 1 µM of AIP-I, a signaling peptide from a different agr group (Williams, Hill, Bonev, & Chan, 2023). Control cultures were grown without AIP-I supplementation. All cultures were grown in anaerobic conditions. As shown in Figure 4A, all strains exhibited similar growth patterns with no notable differences between AIP-I supplemented and unsupplemented conditions. Figure 4B presents fluorescence intensity over time. In the PAG3 strain without AIP-I supplementation, fluorescence intensity increases after approximately 8 hours. In contrast, PAG3 cultures supplemented with AIP-I maintained minimal fluorescence, comparable to the Native strain, which showed consistently low fluorescence under both conditions.

**Figure 4.**
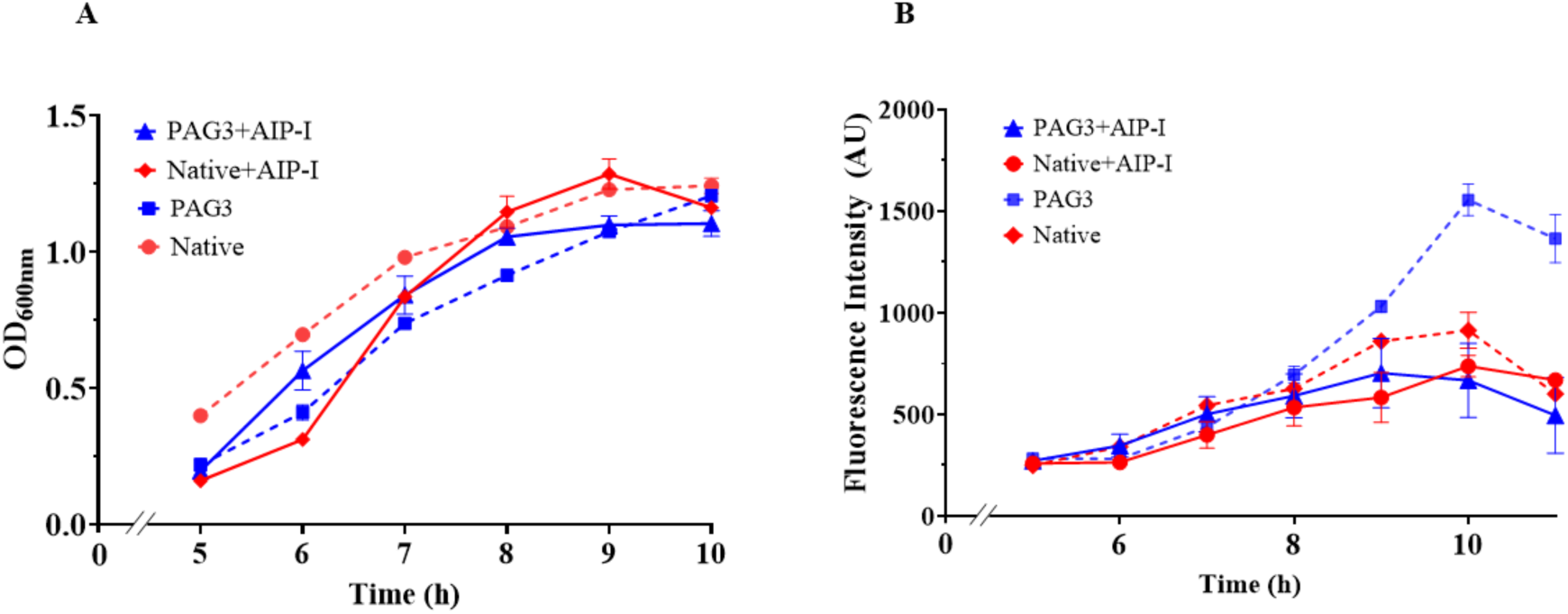
Growth and fluorescence intensity of PAG3 and Native strains with (solid lines) and without (dashed lines) exogenous AIP-I supplementation. (A) OD_600_ growth curves of Native and PAG3 strains cultured in TYG medium, with or without 1 μM AIP-I added at inoculation. (B) Fluorescence intensity over time. An increase in fluorescence is observed in the unsupplemented PAG3 strain, while PAG3 supplemented with AIP-I maintains minimal fluorescence, comparable to the Native strain. The Native strain shows relatively low fluorescence levels in both conditions. Error bars represent one standard deviation from triplicate cultures.

### Engineered agr operon activates at population density threshold

To investigate quorum sensing in the PAG3 strain, we assessed GFP fluorescence intensity in the PAG3 (agr system of *S. aureus* controlling GFP expression), PTG (constitutive GFP expression using the *thl* promoter native to *Clostridium sp.*), and Native (negative control) strains. Cultures were inoculated into anaerobic media and incubated in an anaerobic environment. Measurements of GFP fluorescence intensity and optical density (OD_600_) were initiated 5 hours post-inoculation and continued at intervals until 10 hours. As shown in Figure 5A, the OD_600_ growth curves for PAG3, PTG, and Native strains demonstrate growth. Throughout the growth period, GFP intensity (Figure 5B) in the PTG strain exhibited continual increase, while fluorescence intensity in the Native strain remained relatively constant. The PAG3 strain initially showed low GFP intensity, with values closely matching those of the Native strain; an increase in fluorescence intensity was observed approximately 7 hours post-inoculation. Figure 5C shows fluorescence intensity against population density, showing that the PAG3 fluorescence intensity increases when the bacterial concentration reaches approximately 0.7 OD_600_. As an estimate of per-cell activity, fluorescence intensity per OD_600_ is shown versus cell density in Figure 5D. The PAG3 strain exhibits a considerable relative increase at about 0.7 OD_600_, rising from ∼735 to ∼1885 AU, a ∼2.5-fold increase. In contrast, the intensity remains constant for the Native strain, while the constitutively expressing PTG strain shows a ∼1.4-fold increase (from ∼1800 to ∼2500 AU).

**Figure 5.**
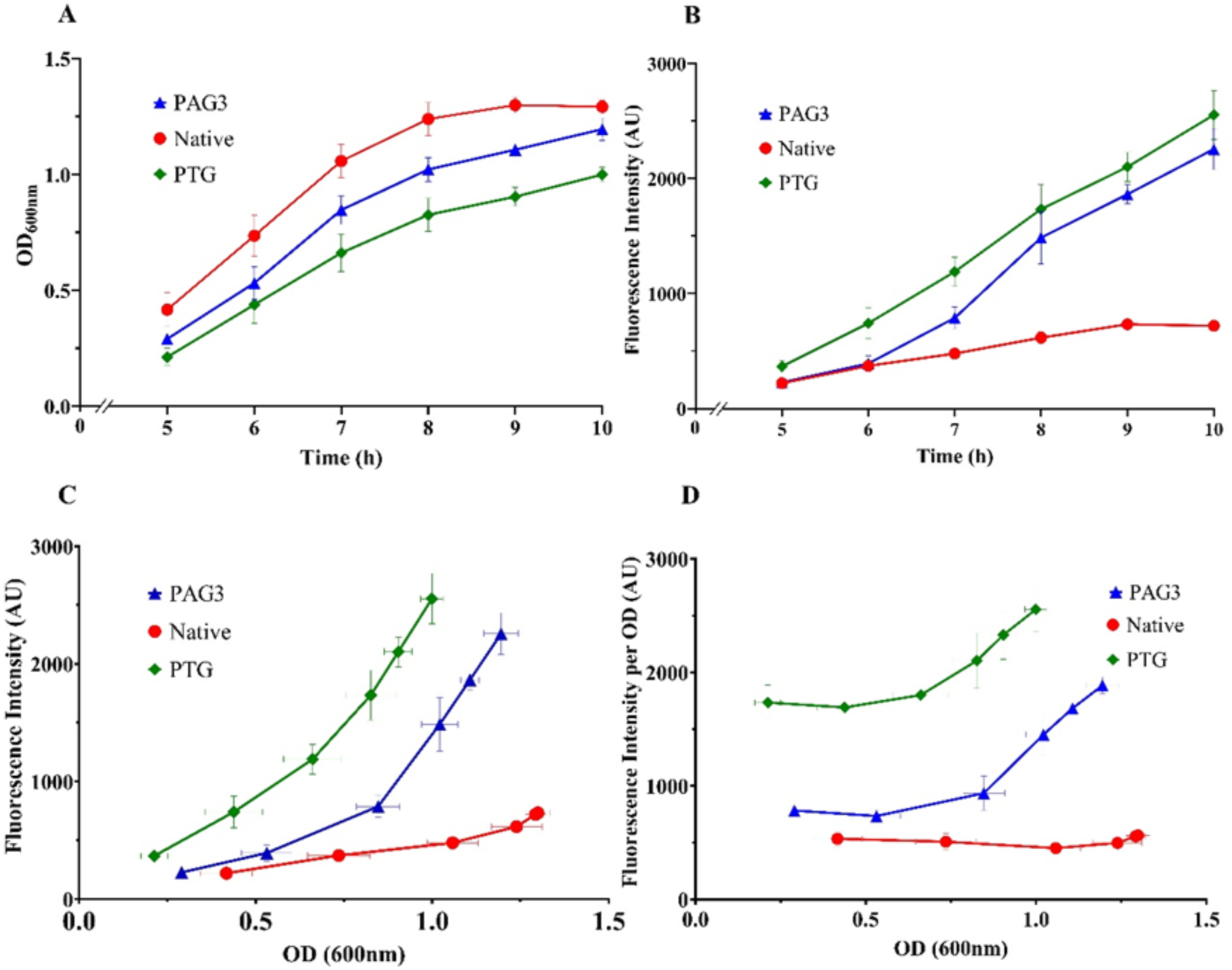
Growth and fluorescence assay of bacteria in batch growth conditions. A) Growth of PAG3, PTG, and Native strains from 5 to 10 hours post-inoculation. B) Fluorescence intensity over time; PAG3 shows fluorescence similar to the Native strain initially, then increases after 6 hours, while Native fluorescence remains constant. PTG, constitutively expressing GFP, shows a continual increase. C) Fluorescence intensity plotted against OD_600_ for PAG3, PTG, and Native strains. The fluorescence of PAG3 at lower OD_600_ is comparable to Native strain, but it increases above OD_600_ ∼0.7. D) Fluorescence intensity normalized by OD_600_ plotted against OD_600_: PAG3 fluorescence is constant at low OD, then increases considerably at OD ∼0.7, while Native fluorescence remains stable. PTG shows an increase as well, but by a relatively modest amount. Error bars indicate one standard deviation from triplicate cultures.

### Removal of accumulated AIP signal delays population density-triggered activation

To further investigate the dependence of QS-induced GFP activity on extracellular AIP concentration, we conducted a growth experiment in which the media, containing accumulated AIP, was replaced mid-way through the growth process. PAG3, PTG, and Native strains were grown in anaerobic TYG media in anaerobic conditions. Fluorescence intensity assays and OD_600_ measurements were initiated 5 hours post-inoculation. At approximately 7 hours post-inoculation, the cells were washed, and media was replaced with new pre-treated TYG media, after which measurements were continued. As a control, separate PAG3 cultures were grown in parallel in batch (no media replacement). As shown in Figure 6A, the OD_600_ growth curve of the Native, PAG3, and PTG strains indicate an increase in bacterial density over time. As expected, the PAG3 culture grown under media replacement reached a higher OD_600_ compared to the batch PAG3.

**Figure 6.**
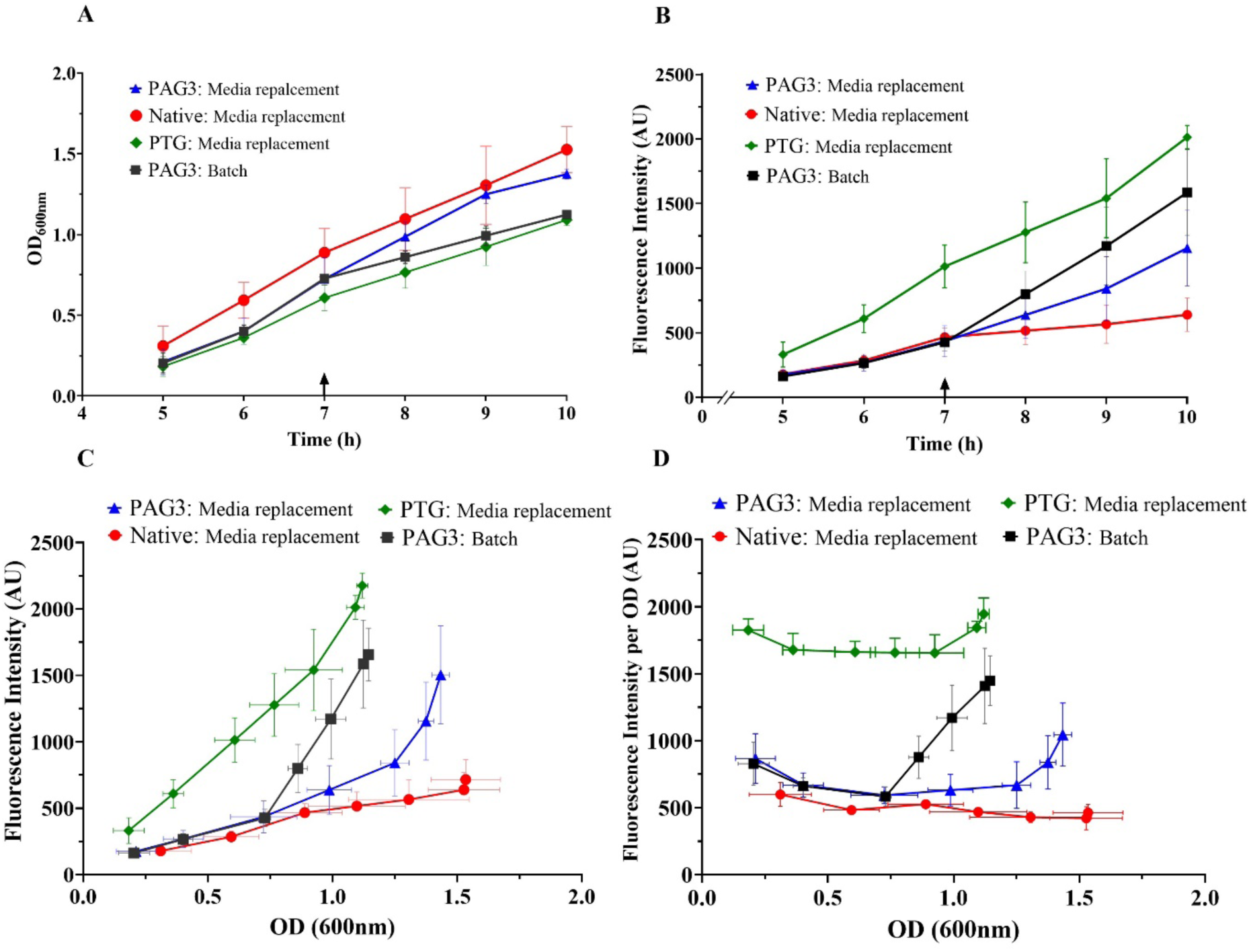
Growth and fluorescence intensity of PAG3, PTG, and Native strains under media replacement conditions. A) OD_600_ growth curves of Native, PAG3, and PTG strains over time, with a control PAG3 strain grown without media replacement (batch). Media replacement at about 7 hours post-inoculation, as indicated (arrow). B) Fluorescence intensity over time, showing a higher increase in batch PAG3 compared to PAG3 with media replacement, a relatively constant fluorescence intensity for the Native, and continual increase for PTG. C) Fluorescence intensity versus bacterial density, indicating that batch PAG3 showed an increase in fluorescence around 0.7 OD_600_, whereas media replacement resulted in fluorescence increase at a higher OD_600_ ∼1.2. D) Normalized fluorescence intensity per OD_600_ versus bacterial density, demonstrating an increase in fluorescence intensity per cell for batch PAG3 at ∼0.7 OD_600_, while media replacement caused the corresponding increase to occur at a higher OD. Normalized fluorescence intensity remained constant for the Native strain and showed a slight increase for PTG. Error bars indicate one standard deviation from triplicate cultures.

As shown in Figure 6B, the initial fluorescence intensity of the PAG3 cultures in both conditions was comparable to that of the Native strain. However, starting about 8 hours post-inoculation, the batch-grown PAG3 showed higher fluorescence than the PAG3 that underwent media replacement. The Native and PTG strains showed expected low and high fluorescence, respectively. Fluorescence intensity activity as a function of bacterial density (OD_600_) is shown in terms of culture fluorescence, in Figure 6C and in terms of fluorescence-per-OD (as a proxy for per-cell fluorescence), in Figure 6D. At low OD (early in the experiment, prior to media replacement), PAG3 showed fluorescence intensity in the same range as the Native strain. The batch-grown PAG3 exhibited a ∼2.5-fold increase in fluorescence intensity as bacterial density progressed from mid-exponential phase to early stationary phase. In contrast, the PAG3 that underwent media replacement shows a delayed increase, achieving about a 1.8-fold increase by the last time-point. As expected, the Native strain did not show a significant increase in fluorescence intensity, and continual increase of fluorescence intensity was observed for PTG.

## Discussion

In the present study, we introduced the *S. aureus* agr quorum sensing (QS) system into *C. sporogenes* to achieve population-dependent gene expression. LC-MS/MS analysis confirmed the production of autoinducing peptide (AIP-III) in the engineered PAG3 strain, with quantification revealing approximately 534 pmol/mL of the biologically relevant isomer after 10 h of anaerobic incubation (Figure 2). No AIP-III signal was detected in the Native strain or blank media. The absence of AIP signal in the Native strain confirms that this specific autoinducing peptide is not produced endogenously by *C. sporogenes*, and that AIP-III production is a result of the introduced *S. aureus* agr system.

To evaluate the specificity and responsiveness of the system, we exposed the engineered strain to both cognate and non-cognate AIP signals. When 1 μM of the *S. aureus* AIP-III was added to culture medium, significant fluorescence intensity was observed in comparison with the unsupplemented control (Figure 3B). Similar responsiveness to exogenous AIPs has been reported in *S. aureus*, where externally supplied AIP restores QS activity in conditions that otherwise suppress agr signaling (Green et al., 2023). Moreover, exogenous AIP has been shown to trigger dispersal of mature *S. aureus* biofilms through agr-mediated protease expression (Boles & Horswill, 2008). In contrast, supplementation with AIP-I, a non-cognate signal peptide, inhibited fluorescence regulated by the engineered agr QS system in PAG3 (Figure 4B), confirming that the engineered system responds specifically to the AIP corresponding to its own agr group. This finding is consistent with the well-established cross-inhibition behavior of *S. aureus* agr groups, where non-cognate AIPs act as antagonists by binding AgrC without activating downstream signaling: AIP-I from agr group I does not activate the AgrC receptor from group III and instead competitively inhibits its response (Le & Otto, 2015; Williams et al., 2023). The lack of response to AIP-I in our engineered strain mirrors these native specificities and further demonstrates the precision of the introduced circuit. Together, these results confirm that the synthetic agr system retains both functionality and signal specificity in *C. sporogenes*, with gene expression modulated effectively by cognate AIP accumulation or supplementation.

We assessed the behaviour of the engineered QS system in culture growth by measuring agr-activated GFP fluorescence intensity. The engineered PAG3 strain exhibited distinct fluorescence intensity patterns compared to the Native strain (Figure 5). When colonies of PAG3, PTG, and Native strains were cultured in anaerobic conditions, density-normalized fluorescence intensity of PAG3 was initially at the same level as that of the Native strain. In contrast, PTG exhibited consistently higher density-normalized fluorescence intensity. At about 7 hours post-inoculation, when its OD_600_ reached approximately 0.7, the density-normalized fluorescence intensity of PAG3 began to increase, while the Native strain’s remained constant. This behaviour suggests that the engineered agr operon in PAG3 activates gene expression once the bacterial density reaches this threshold.

To further probe the response of the QS system to AIP accumulation, we conducted a media replacement experiment. We hypothesized that refreshing the media during the early exponential phase would remove accumulated AIP-III, effectively interrupting the quorum sensing process by ‘resetting’ the AIP level. Following media refreshment, the increase in density-normalized fluorescence intensity of PAG3 occurred later and at higher bacterial density compared with PAG3 grown without media replacement (Figure 6). This finding aligns with previous studies demonstrating that QS in Gram positive bacteria relies on AIP accumulation to activate QS-regulated gene expression (Elisabeth, Jamie, P., & Klaus, 2012; Novick & Geisinger, 2008; Williams et al., 2023). The observed delay in the response following media replacement is consistent with studies showing that alterations in the extracellular environment can interfere with Gram positive QS activation dynamics (Kavanaugh & Horswill, 2016; Williams et al., 2023). Previous studies have demonstrated that changes in extracellular conditions, including nutrient fluctuations and removal of accumulated signaling molecules, impact the timing and strength of QS responses (Horswill, Stoodley, Stewart, & Parsek, 2007).

Turning to potential applications of this system, we note that QS-based synthetic bacterial designs have enabled precise, density-dependent regulation of therapeutic molecule production, facilitating controlled drug release and synchronized bacterial behavior in pathological environments (Dang et al., 2023; Hwang, 2024). While earlier studies employed QS in organisms such as *E. coli* and *Salmonella* to develop QS-controlled therapeutic systems for tumor targeting (Chowdhury et al., 2019; Din et al., 2016; Swofford et al., 2015), our work extends these principles to *C. sporogenes*, a clinically promising but genetically less tractable obligate anaerobe. As introduced above, *Clostridia*, particularly *C. sporogenes* (formerly classified as *C. butyricum* M55 or *C. oncolyticum* M55 (Minton, 2003)), have emerged as promising agents for cancer therapy due to non-pathogenicity (Kubiak et al., 2015) and selective germination of spores in hypoxic tumor regions. In addition, the relatively large genome size and coding capacity of *C. sporogenes,* compared to other proteolytic strains, such as *C. novyi*-NT (an attenuated strain of *Clostridium novyi* (M. Zygouropoulou, A. Kubiak, A. V. Patterson, 2019)), make it an attractive chassis for synthetic biology applications aimed at enhancing tumor therapeutic delivery. *C. sporogenes* has shown significant potential in tumor regression, demonstrating efficient tumor colonization (1– 2×10^8^ vegetative bacteria/g of tumor (Liu et al., 2002) and inherent oncolytic capabilities (Bhave, Hassanbhai, Anand, Qian Luo, & Teoh, 2015). Upon intravenous injection, spores of *C. sporogenes* selectively colonize necrotic and hypoxic regions of solid tumors, where they produce degradative enzymes that contribute to tumor lysis (M. Zygouropoulou, A. Kubiak, A. V. Patterson, 2019). Additionally, *C. sporogenes* can be genetically modified to serve as a targeted delivery vehicle for therapeutic agents, offering a bacteria-mediated approach to localized drug delivery (Kubiak, Bailey, Dubois, Theys, & Lambin, 2021; Theys et al., 2006). The engineered QS system in *C. sporogenes* described here offers the potential to provide tunable, population-dependent control over gene expression, ensuring that bacterial therapies activate only within tumor environments at optimal bacterial densities to maximize efficacy while minimizing off-target effects.

Quorum sensing (QS) in *Clostridia* species also has potential in biomanufacturing, where these bacteria are widely used for industrial fermentation processes, including acetone-butanol-ethanol (ABE) fermentation (focused on *C. acetobutylicum*) and the biosynthesis of various chemicals and fuels (Xue, Zhao, Chen, Yang, & Bai, 2017). Focusing specifically on *C. sporogenes*, this strain has been studied for non-acetone butanol production from rice straw (Gottumukkala et al., 2013, 2015) and has been identified as a producer of 3-indolepropionic acid (IPA), a potent antioxidant with neuroprotective properties (Du, Qi, Wang, Liu, & Wu, 2021). Engineered QS circuits in *Clostridia* species could be used to optimize solventogenesis, metabolite flux, and stress tolerance, thereby improving biofuel yields. Additionally, QS-controlled regulation of sporulation and stress resistance genes could enhance fermentation stability, prolonging microbial viability in industrial bioreactors (Cheng, Bao, & Yang, 2019; Papoutsakis, 2008). Leveraging synthetic QS systems in *Clostridia* represents a promising strategy for increasing biofuel production efficacy and sustainability in industrial settings.

## ASSOCIATED CONTENT

### Supporting Information

The Supporting Information is available free of charge at the ACS Publications website. – Primer sequences and additional figures referenced in the main text (PDF)

## AUTHOR INFORMATION

### Author Contributions

Sara Sadr: Conceptualization, formal analysis, investigation, methodology, writing – original draft, and writing – review & editing. Bahram Zargar: Conceptualization, formal analysis, funding acquisition, investigation, project administration, resources, supervision, and writing – review & editing. Marc Aucoin: Formal analysis, investigation, project administration, supervision, and writing – review & editing. Brian Ingalls: Conceptualization, formal analysis, funding acquisition, investigation, project administration, resources, supervision, and writing – review & editing.

## ACKNOWLEDGMENT

We thank Dr. Farzad Kobarfard for his valuable support and technical expertise in LC-MS/MS analysis. We also gratefully acknowledge Dr. Matthew LaBore for his guidance and assistance in the molecular biology aspects of this work, particularly during plasmid construction and development.

