## Supplement for "Construction and Functional Characterization of a Heterologous Quorum Sensing Circuit in *Clostridium sporogenes*"

**Supplementary Information**

**A**

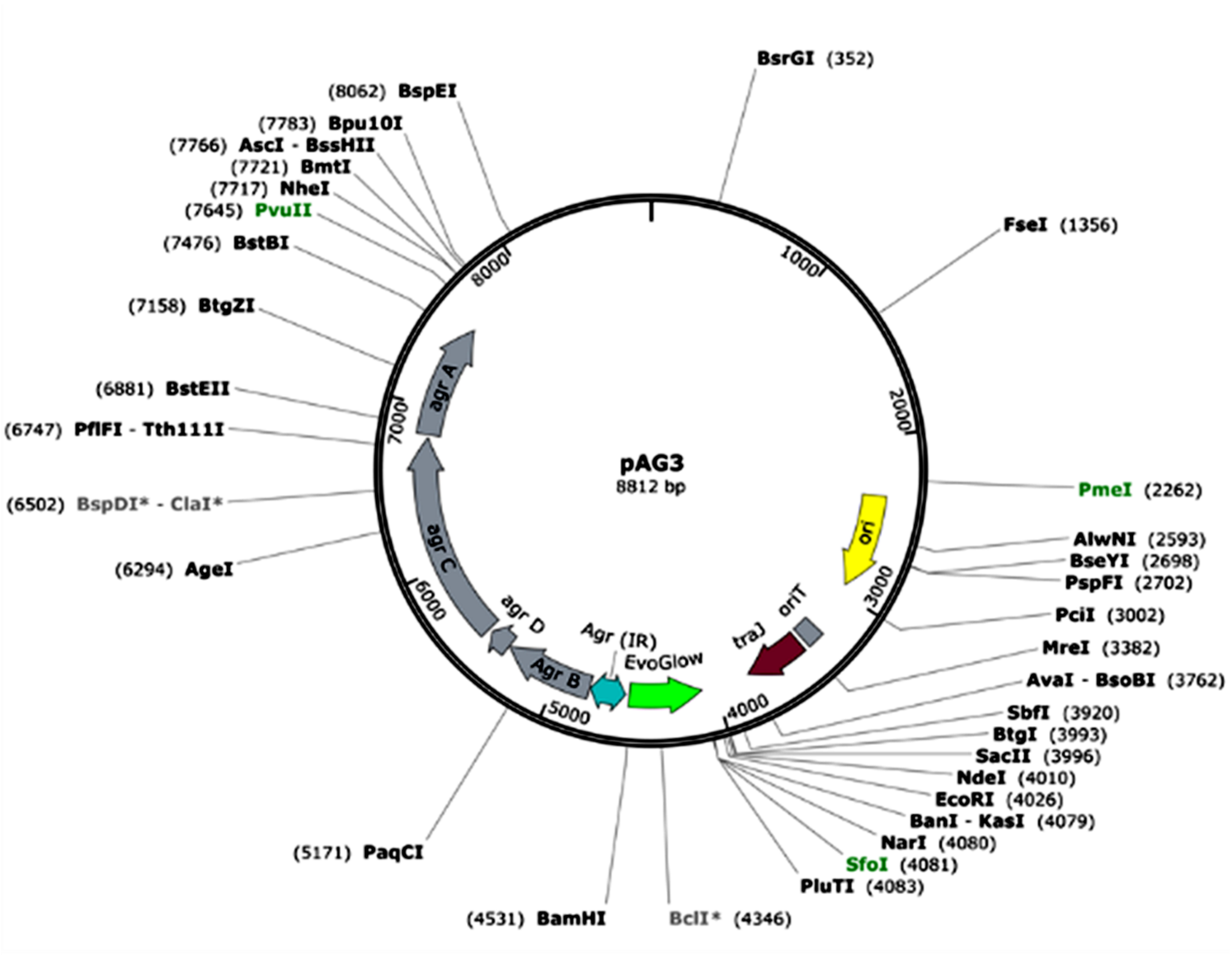

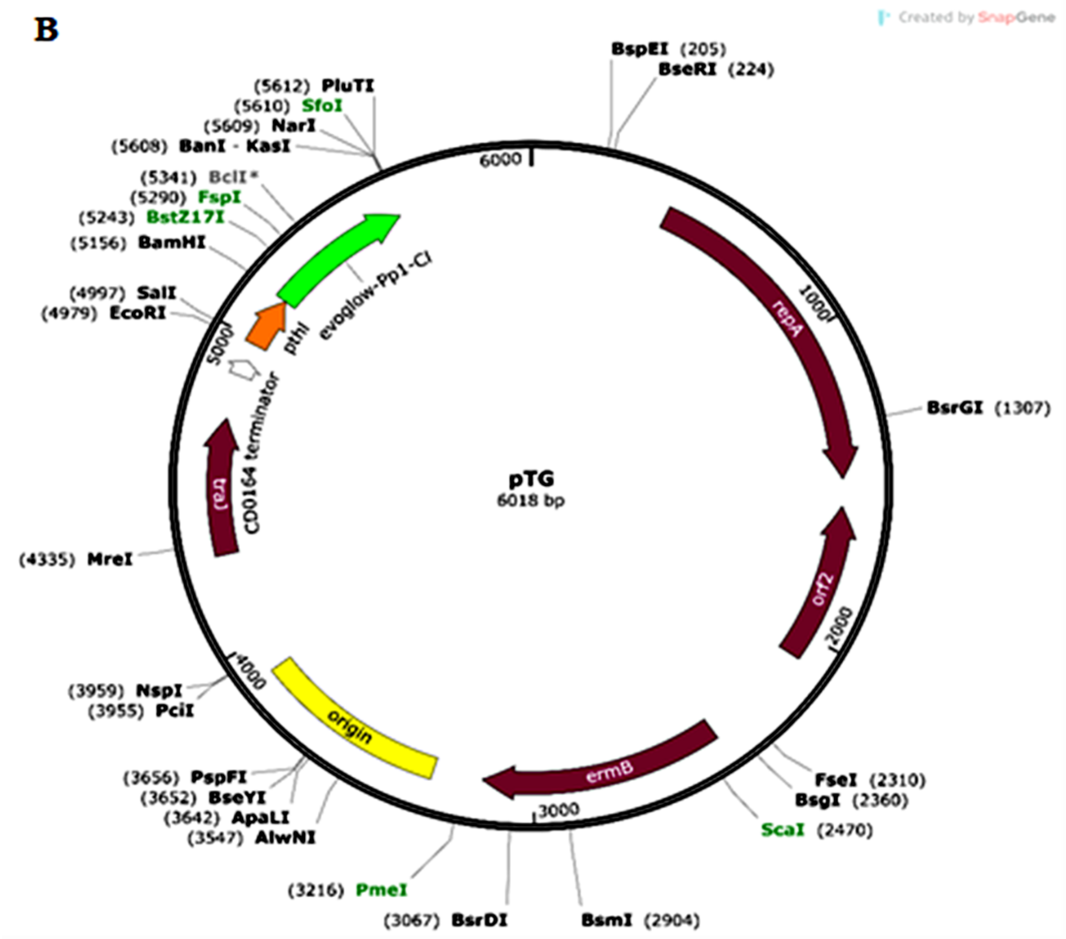


**Figure S1.** Plasmid maps of pAG3 (A), pTG (B). The PAG3 strain was constructed by transformation of native *C. sporogenes* with the pAG3 plasmid. The plasmids of pTG was transformed into *C. sporogenes* to develop the control strains of PTG.

**Table S1-** Primers used in PCR to construct the plasmids

| Primer | Sequence |
|---|---|
| pglow_f_XbaI | GGCATTCTAGAGTAGGATCAAGGAGGTTAGTTAGAATGGG |
| pglow_r_pstI_spei | CTGCAGCGGCCGCTACTAGTCTGGCAAATCATTAAGTGGCGCCTT |
| Agr43300_fwd | GTTAACTGACTTTATTATCTTA |
| Agr43300_rev | TTCCATCACATCTCTGTGATCT |
| thl_F | GGCATGAATTCGGTTGGAATGGCGTGTGTGTTAGCCA |
| Agr_fwd | CCTTGATCCTACTTCCATCACATCTCTGTGATCTAG |
| Agr_rev | GGCAGCCGATCTTATTATATTTTTTTAACGTTTCTCACCG |
| P_fwd | AATAAGATCGGCTGCCCGTCGTTTTACAACGTCGTG |
| P_rev | TGTGATGGAAGTAGGATCAAGGAGGTTAGTTAGAATGGG |
